# Synthetic communities of maize root bacteria interact and redirect benzoxazinoid metabolization

**DOI:** 10.1101/2025.03.12.642850

**Authors:** Lisa Thoenen, Tobias Zuest, Christine Pestalozzi, Marco Kreuzer, Pierre Mateo, Gabriel Deslandes, Christelle A.M. Robert, Rémy Bruggmann, Matthias Erb, Klaus Schlaeppi

## Abstract

Plant roots are colonised by diverse microbial communities. These communities are shaped by root exudates including plant specialized metabolites. Benzoxazinoids are such secreted compounds of maize. Individual microbes differ in their ability to tolerate and metabolize antimicrobial benzoxazinoids. To investigate how these traits combine in a community, we designed two synthetic communities of maize root bacteria that share six common strains and differ in their ability to metabolize benzoxazinoids based on the seventh strain. We exposed both communities to the benzoxazinoid MBOA (6-methoxybenzoxazolin-2(3H)-one) and found that the metabolizing community did not degrade MBOA to its aminophenoxazinone, as observed for individual strains, but as a community they formed the corresponding acetamide. MBOA shaped differential compositions of both communities and increased the fraction of MBOA-tolerant strains. The benzoxazinoid metabolizing community showed a higher tolerance to MBOA and was able to utilize MBOA as their sole carbon source for growth. Hence, bacterial interaction results in alternative benzoxazinoid metabolization and increases community performance in presence of these antimicrobial compounds. Future work is needed to uncover the genetics of this metabolic interaction and ecological consequences for the bacterial community and the host plant.

**Importance:** We investigated how maize root bacteria - alone or in community - tolerate and metabolize antimicrobial compounds of their host plant. We found the capacity to metabolize such a compound to impact bacterial community size and structure and most importantly, to benefit community fitness. We also found that interacting bacteria redirected the metabolisation of the antimicrobial compound to an alternative degradation product. Our work highlights the need to study the team work of microbes to uncover their community traits to ultimately understand the ecological consequences for the bacterial community and eventually the host plant.

## Introduction

Most multicellular organisms are closely associated with microbial communities, which provide important functions to the host. Plant microbiomes function in promoting growth, providing nutrients, protecting from disease but some microbiome members can also act as pathogens (1–3). The microbial members including bacteria, fungi, protists and oomycetes co-adapted with their host over long evolutionary time scales (4–7). Root exudates, which can account for up to one-fifth of the plant’s assimilated carbon (8) contain primary metabolites including sugars, amino acids, organic acids and fatty acids, as well as secondary metabolites. The latter, also called specialized metabolites, act in diverse ways to govern the interactions with the plant’s environment. Often these compounds have antimicrobial function, selecting adapted microbes and thereby shaping the species-specific root microbiome (8–12). Examples include glucosinolates, camalexins, triterpenes, and coumarins in *Arabidopsis thaliana* (9), the saponin tomatine from tomato (13), and benzoxazinoids (14–18), diterpenoids (19), zealexins (20) and elavonoids (21) in maize. While there is increasing knowledge how individual microbes react to these compounds, it is largely unknown how they cope as communities with antimicrobial root exudates.

Microbial community members not only co-evolved with their hosts but they also adapted to interact efficiently with each other. On one hand they developed mechanisms to optimize their fitness through competitive mechanisms such as the production of antimicrobial compounds (22). On the other hand, some also adapted to profit from each other to efficiently use resources and accomplish tasks which are not feasible as individuals (23, 24). For example, community members might exchange metabolites to complement each other’s biosynthetic pathways (25). Such interactions cannot only improve community fitness but also alter final product distribution, and give rise to new phenotypes without genetic modification (26). Microbes often cooperate to degrade antimicrobials or pollutants (24, 27) or metabolize complex substrates (28). Examples include the metabolization of the herbicide atrazine by an adapted four-strain consortium in which one bacterium metabolized the herbicide to cyanuric acid, which then served as a carbon source for the other community members (29). Or, another three-strain consortium converted the herbicide linuron more efficiently in co-culture showing increased expression of the linuron hydrolase (30). Also, the degradation of antibiotics by one community member affected the community composition (31). While there are examples of how microbial communities cooperate to degrade antimicrobial compounds and pollutants, little is known about how root microbiome members interact to cope with antimicrobial plant metabolites.

Crops belonging to the family of sweet grasses (*Poaceae*) include maize, wheat and rye produce benzoxazinoids as specialized metabolites (32, 33). These indole-derived alkaloids mainly function in protecting the plant from insect pests and pathogens (32, 34, 35), providing support in iron uptake (36) and structuring the root microbiomes by acting as selective antimicrobials (11, 14–18). **Fig. S1** contains the full names and structures of all abbreviated compounds mentioned in this study. DIMBOA-Glc is the main root-exuded benzoxazinoid of maize (16), and its chemical fate in soil is well understood: Upon exudation, DIMBOA-Glc is deglycosylated by microbe- and/or plant-derived enzymes (35) to form DIMBOA, which spontaneously converts to more stable MBOA (37). In soil, MBOA has a half-life of a few days and, when further metabolized by microbes to reactive aminophenols (35), MBOA can take three different routes: route (I) is favoured under aerobic conditions forming aminophenoxazinones such as AMPO through oxidation (38); route (II) results in acetamides such as HMPAA through acetylation (39), or alternatively, route (III) yields malonic acids such as HMPMA through acylation (39). Route I is probably most relevant for the rhizosphere, and the resulting AMPO remains detectable in the soil of maize fields for up to a few months (37). Several bacterial isolates were shown to metabolize MBOA to AMPO, which is mediated by the lactonase BxdA (40). Route II was demonstrated for the soil fungus *Fusarium sambucus*, which metabolizes BOA to the acetamide HMPAA (39). Following Route III, the maize seed endophytic fungus *Fusarium verticillioides* converts BOA to the malonamic acid HPMA catalysed by a metallo-β-lactamase (41). There is first evidence that benzoxazinoid metabolization routes vary when different microbes interact. The fungus *Fusarium verticillioides* was co-cultured with the bacterium *Bacillus mojavensis* in presence of BOA and metabolization was redirected from HMPAA to the aminophenoxazinone APO (42). Hence benzoxazinoid metabolization in complex microbial communities warrants further research, for instance to test if microbes metabolically redirect benzoxazinoid degradation.

To study the mechanisms governing composition and functions of microbial communities and their interactions with the host or the environment, synthetic communities (SynComs) are a useful tool (43). A SynCom is a microbial community created by rationally mixing strains from a collection from a specific environment. Strains can be added, eliminated, substituted or individual strain functions manipulated, thereby allowing the detailed study of SynCom members, their functions and the resulting community performance (45). Although, these reduced systems do not accurately represent nature, they allow the replication of phenotypes mediated by the microbiome.

Here we aimed to test if maize root bacteria interact to tolerate and metabolize the benzoxazinoid MBOA in small SynComs. We investigated I) the type of degradation metabolites produced by SynComs and which SynCom members are responsible for metabolization, II) how the ability to metabolize benzoxazinoids affects community growth and tolerance, and III) if MBOA shapes the composition of the two SynComs differently depending on their ability to metabolize MBOA. To address these questions, we designed two SynComs consisting of six common core strains and one variable *Microbacterium* strain that differed in the ability to metabolize MBOA. We exposed the SynComs to MBOA and measured community growth, community tolerance, community composition and benzoxazinoid metabolite profiles. We found the SynCom containing the benzoxazinoid metabolizing *Microbacterium* to convert MBOA to HMPAA, to grow on MBOA as a sole carbon source and to be more tolerant to MBOA. Interestingly, the benzoxazinoid metabolite HMPAA was not formed by single strains but only in the SynCom in combination of *Microbacterium* LMB2 with another strain. Our results revealed that maize root bacteria interact to metabolize and tolerate benzoxazinoids, which is beneficial for microbial growth.

## Results

### In community, the bacteria metabolize MBOA to HMPAA instead of AMPO

In previous work, we characterized individual maize root bacteria for their abilities to tolerate (11) and metabolize benzoxazinoids (40). In this study, we investigated how these traits function in a community context. To this end we examined growth and composition of a synthetic community (SynCom) of maize root bacteria and the metabolites they produce after exposure to MBOA. We constructed two 7-member SynComs consisting of taxonomically distinct bacteria (distinguishable by 16S rRNA gene amplicon sequencing) that differed in their tolerance to MBOA (**Fig. 1A, Fig. S2**). Both SynComs shared a common core of six bacteria that do not degrade MBOA in liquid culture (see notes on LMX9231 in **Supplementary Results**, **Fig. S3**). To complete the communities, alternative *Microbacterium* strain, differing in their ability to metabolize MBOA, were added to each. This resulted in a *non*-metabolizing SynCom (nonSC) containing *Microbacterium* LMI1x which is unable to metabolize MBOA, and a MBOA-*metabolizing* SynCom (metSC) containing the MBOA degrader *Microbacterium* LMB2 (**Fig. 1A**). Importantly, monocultures of *Microbacterium* LMB2 produce primarily AMPO after exposure to MBOA (**Fig. S3B**).

**Figure 1:**
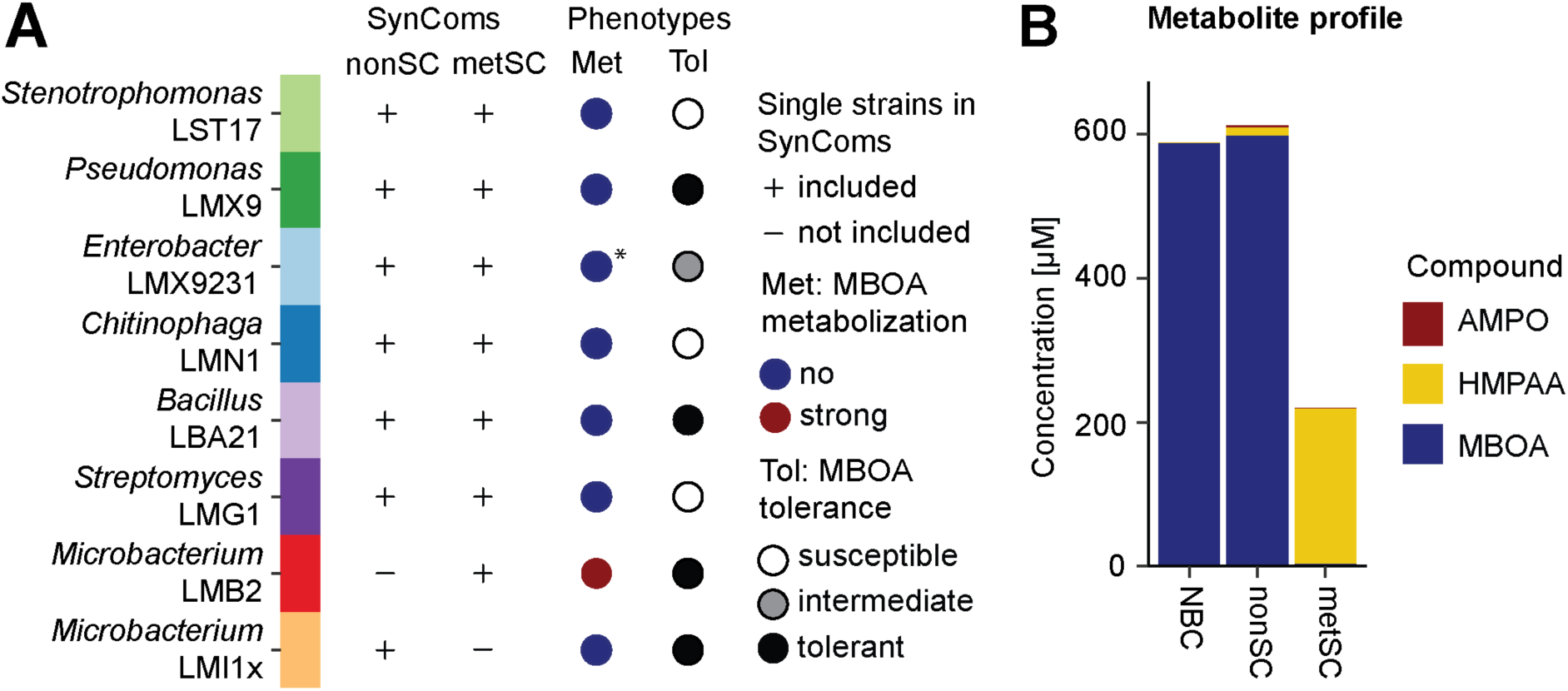
SynCom design and metabolisation of MBOA. **A)** List of bacterial strains selected for the non-metabolizing SynCom (nonSC) and the metabolizing SynCom (metSC) indicated with a “+” if included and a “-“ if not included. The phenotypes of each strain in MBOA metabolization (Met, data from Ref 40) and MBOA tolerance (Tol, based on 2’500 µM data from **Fig. S2**) are listed in columns three and four. (*) LMX9231 does not metabolize MBOA to AMPO in liquid culture, see notes in Supplementary Results. **B)** The nonSC and metSC SynComs were grown in 96-well plates containing 50% TSB supplemented with 500 µM MBOA for 68 h and afterwards the metabolisation of MBOA to AMPO and/or HMPAA was determined. SynComs and their no bacteria controls (NBC, media with MBOA but without bacteria) were grown in triplicates and pooled before measurements (n = 1).

To test community effects, we cultured the nonSC and metabolizing metSC in 96-well plates in complex medium supplemented with 500 µM MBOA. We performed metabolite analyses to confirm their differential MBOA degradation and to determine the various metabolization products at 68 h (**Fig. S1, see Materials and Methods**). We found that the nonSC did not degrade MBOA as expected whereas the metSC degraded MBOA and formed HMPAA rather than AMPO (**Fig. 1B**). Over the course of the experiment, the levels of MBOA continuously decreased, HMPAA continuously increased and not much AMPO accumulated (**Fig. S4A**). In addition, we also detected a metabolite feature with a mass corresponding to the intermediate AMP (acetylation of AMP yields HMPAA; **Fig. S1**) that accumulated over time similar to HMPAA (**Fig. S4B**). In summary, the two SynComs nonSC and metSC differ in their ability to metabolize MBOA. In contrast to pure cultures of LMB2 that primarily produce AMPO (**Fig. S3B**), we measured predominantly the MBOA metabolization product HMPAA in the metSC where LMB2 is present in combination with other strains. This finding indicates that *Microbacterium* LMB2 interacts with at least one other metSC strain, resulting in the formation of HMPAA.

### Most strains can form HMPAA together with LMB2

To identify the strain(s) that are involved in HMPAA formation, we first tested reduced metSC communities in which individual strains were dropped out. We incubated these ‘dropout SynComs’ in complex medium supplemented with 500 µM MBOA for 68 h and performed metabolite analyses to determine the various metabolization products (**Fig. S1**). First, the dropout SynCom without *Microbacterium* LMB2 did not degrade MBOA (**Fig. 2A**), confirming the key role of LMB2 in initiating the metabolization of MBOA. Second, HMPAA was formed in all other dropout SynComs in which another strain was removed. While the dropout of *Enterobacter* LMX9231 and *Stenotrophomonas* LST17 did not markedly alter the metabolizing capacity of the metSC, considerable levels of MBOA were still detected in the dropout SynComs lacking any of the other four strains. Overall, this indicated that not one single specific strain is required for formation of HMPAA in the metSC.

**Figure 2:**
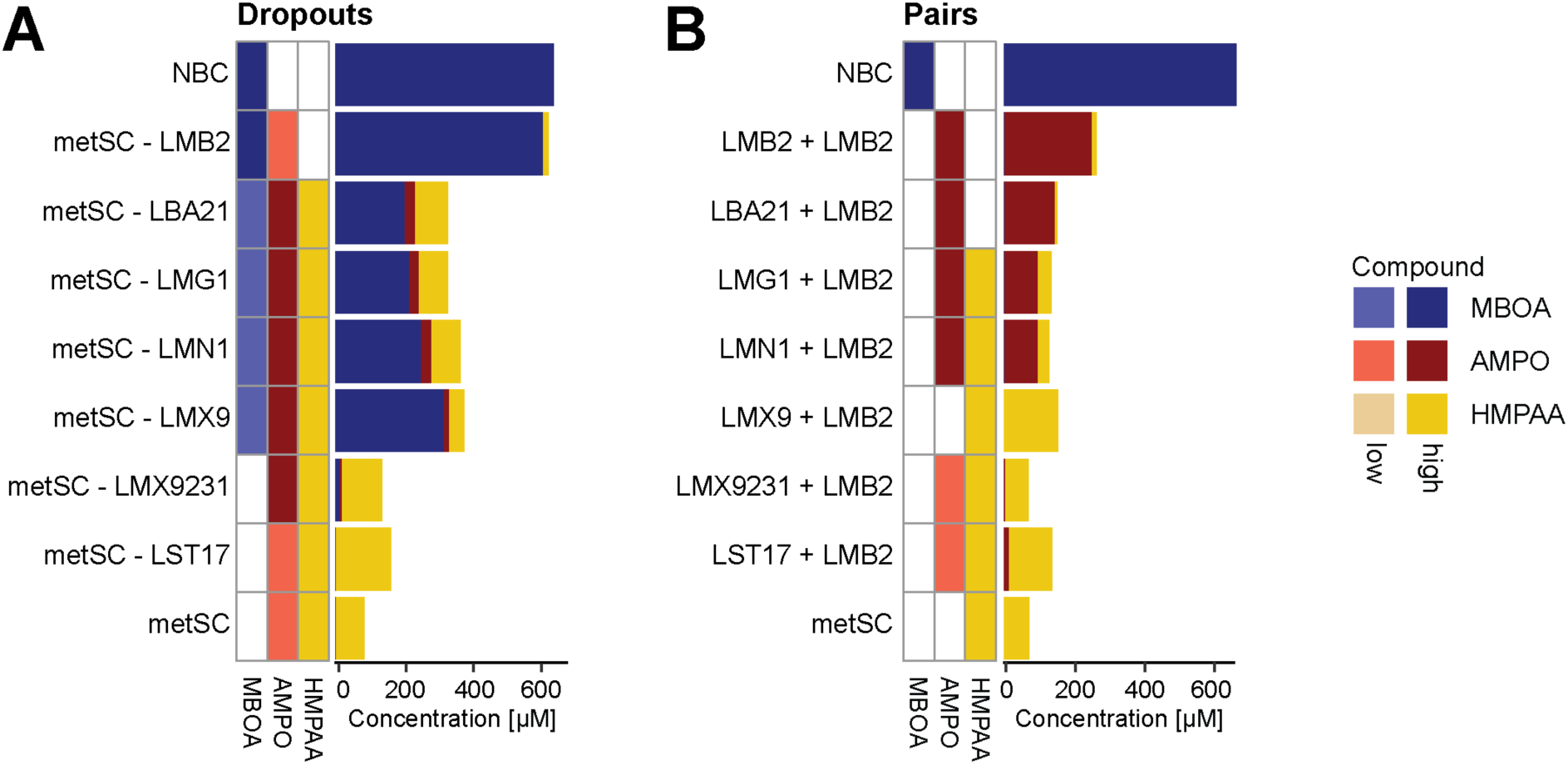
Metabolite proKiles. MBOA, AMPO and HMPAA were measured for dropout SynComs (A) and paired cultures (B) after growth with MBOA. Heatmaps display the qualitative detections of MBOA, AMPO and HMPAA. Detected compound levels were classified as “low” or “high” and are indicated with light and dark color tones, respectively. Stacked bargraphs display the quantitative metabolite measurements. SynComs and paired strains were grown in 96-well plates in liquid 50% TSB supplemented with 500 µM of MBOA alongside their no bacteria controls (NBC, media with MBOA but without bacteria). Cultures were grown in triplicates and pooled before measurements (n = 1).

For further identification of individual strains interacting with *Microbacterium* LMB2 to form HMPAA, we tested each community member paired with LMB2 as in the assay above. Consistent with previous work (33; **Fig. S3B**), pure cultures of *Microbacterium* LMB2 completely degraded MBOA and formed primarily AMPO (**Fig. 2B**). In co-cultures with *Bacillus* LBA21, *Streptomyces* LMG1 and *Chitinophaga* LMN1, AMPO remained the primary product measured. In contrast, in co-cultures with *Pseudomonas* LMX9, *Enterobacter* LMX9231 and *Stenotrophomonas* LST17 little or no AMPO was detected, similar to the full metSC. Importantly, with the exception of the co-culture with *Bacillus* LBA21, most pairs formed HMPAA. The highest levels of HMPAA were detected in co-cultures with *Pseudomonas* LMX9 and *Stenotrophomonas* LST17, intermediate levels with *Enterobacter* LMX9231 and the full metSC and low levels with *Streptomyces* LMG1 and *Chitinophaga* LMN1. Taken together, HMPAA can be formed with five out of the six core members of the metSC in co-culture with *Microbacterium* LMB2. According to the chemical model (**Fig. S1**), we hypothesize that *Microbacterium* LMB2 initiates the metabolization of MBOA by opening its lactone ring by BxdA (40) and forming the intermediate AMP, which can then be further acetylated to HMPAA by each of these five strains. In absence of these strains, the intermediate AMP undergoes oxidative dimerization to AMPO.

### The metabolizing SynCom exhibits growth in the presence of MBOA

Next, we tested whether the ability to metabolize MBOA enables the metSC to use MBOA as a sole carbon source for bacterial growth. For this we grew the nonSC and metSC SynComs in minimal media supplemented with 500 or 2’500 µM MBOA, glucose (Glc) or no additional carbon source (DMSO), keeping the DMSO concentration between treatments constant. We quantified community growth by monitoring optical density (OD_600_) over time. While both SynComs exhibited similar growth in the DMSO and glucose controls, the metSC grew significantly better in MBOA than the nonSC, particularly at the high MBOA concentration (**Fig. 3**). Testing all strains individually confirmed that *Microbacterium* LMB2 was the only strain capable of utilizing MBOA as a sole carbon source (**Fig. S5**). This analysis suggested that the metabolizing SynCom metSC benefitted from LMB2’s capacity to use MBOA as a sole carbon source. Of note, this conclusion comes with the caveat that the OD_600_ measurements do not discriminate between the growth of LMB2 and the other community members (see below).

**Figure 3:**
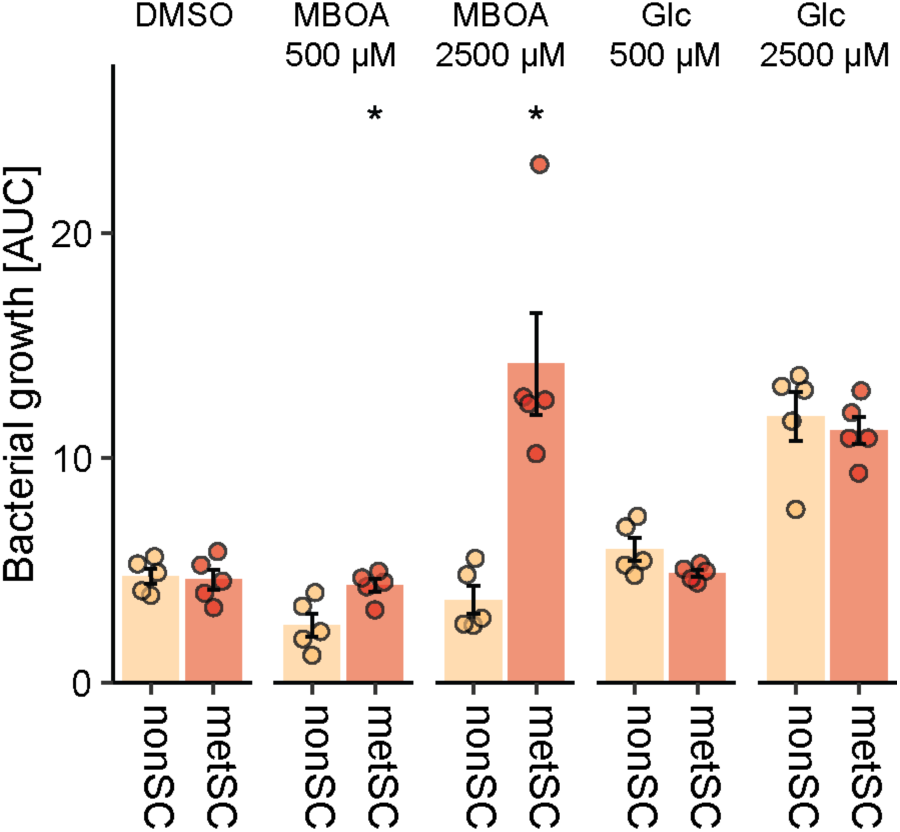
The metSC SynCom can use MBOA as sole carbon source. The two SynComs were grown in 96-well plates in minimal medium supplemented with 500 and 2’500 µM MBOA or glucose (positive controls). Minimal medium supplemented with DMSO only was used as negative control. Bacterial growth over 68 h is reported as the area under the growth curve (AUC) derived from OD_600_ measurements over time. Means ± standard error and individual data points are shown (n = 5). Asterisks indicate significant differences between treatment (pairwise t-test, Bonferroni-adjusted P < 0.05).

### The capacity to metabolize MBOA affects community size and structure

To examine the effect of the MBOA metabolization trait on community size, we next exposed both SynComs in complex medium to 500 or 2’500 µM of MBOA or a control treatment (DMSO) for 68 h. For scale and sampling reasons we performed this experiment in Erlenmeyer flasks and first confirmed that MBOA was degraded by the metSC but not by the nonSC in this experimental setting and that HMPAA is also the dominant degradation product under these conditions (**Fig. S6A&B**). We found that the total community size, as determined by colony-forming units (CFU), decreased with increasing levels of MBOA but that the metSC showed a larger community size than the nonSC in presence of MBOA (**Fig. 4A**). Complementary, we also determined the OD_600_ of the cultures and found similar results (**Fig. S6C**), with both CFU- and OD-based community quantifications correlating well (**Fig. S6D**). This revealed that the ability to metabolize MBOA enabled the metSC community to reach a higher population size when faced with the antimicrobial compound.

**Figure 4:**
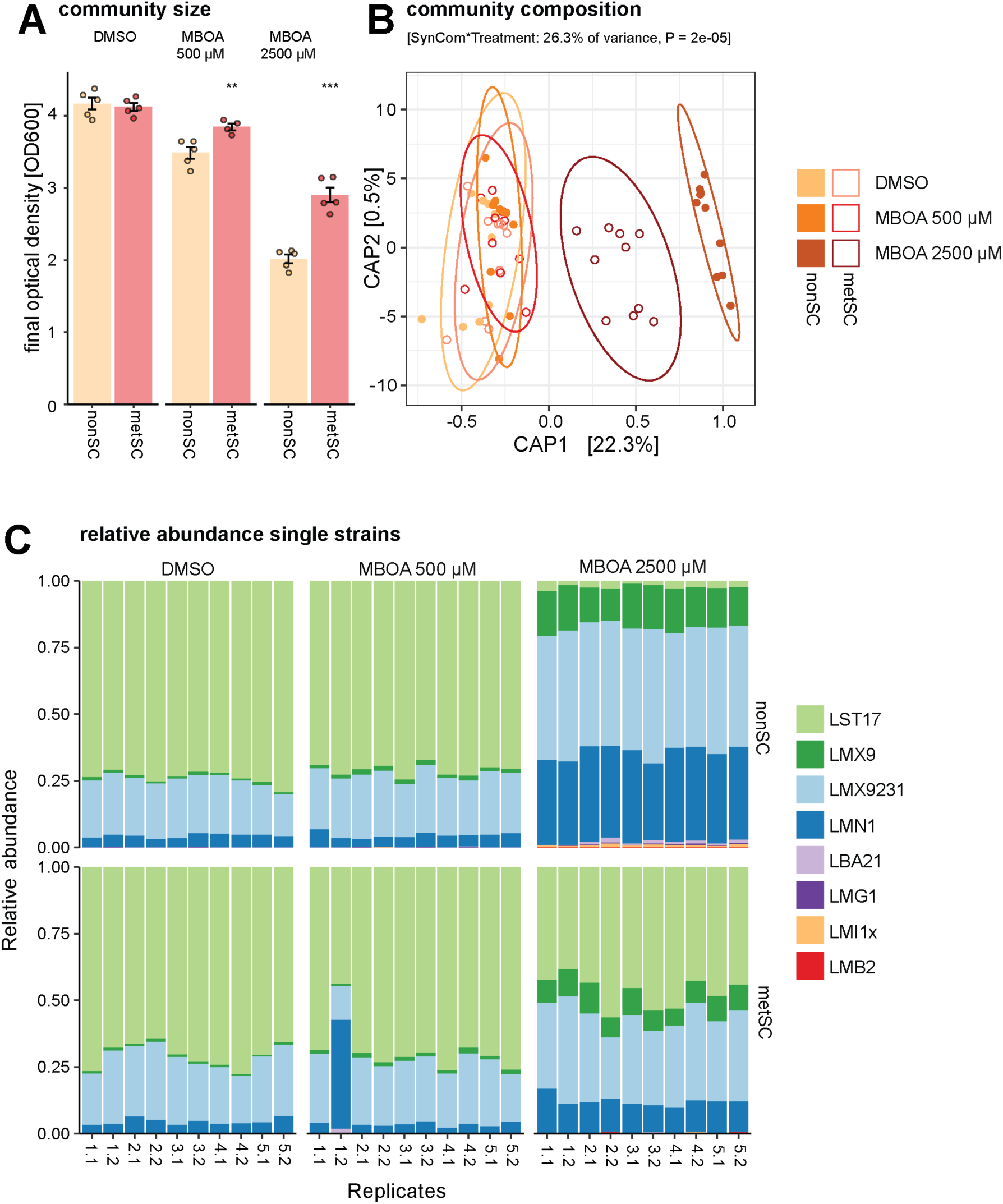
Size and composition of the nonSC and metSC SynComs in presence and absence of MBOA. SynComs were grown in shake flasks in 50% TSB supplemented with 500 µM or 2500 µM MBOA or DMSO only for 68 h. Five replicate cultures were set up for each SynCom and treatment and two parallel samples of each culture were collected for community analysis by 16S rRNA gene amplicon sequencing. **A)** Bacterial growth determined by OD_600_ at the end of the experiment (n = 5). Asterisks indicate significant differences between the two SynComs (pairwise t-test, Bonferroni-adjusted P < 0.05) **B** and **C)** Community compositions at the end of the experiment. Shown are a Canonical analysis of Principal Coordinates (CAP) based on Bray-Curtis distances and the model ∼SynCom * treatment (n = 10) **(B)** and relative abundances of the SynCom members for individual samples **(C)**.

In the same experiment, we also determined community compositions of the two SynComs using amplicon sequencing. While the presence of MBOA significantly affected community composition (PERMANOVA, R^2^ = 0.26, *P* = 0.001), no major effect was found for the two types of SynComs or their interaction with MBOA (**Table S1**). Constrained ordination visualized this MBOA effect along the first axis with SynCom compositions differing at the high MBOA concentration relative to communities at 500 µM MBOA and the DMSO controls (**Fig. 4B**). The high MBOA treatment (2’500 µM) had a stronger impact on the composition of the nonSC than on the metSC. In DMSO and at 500 µM of MBOA, both SynComs were compositionally dominated by *Stenotrophomonas* LST17, followed by lower levels of *Enterobacter* LMX9231 and *Chitinophaga* LMN1 while the other strains were either only detected at very low levels or not any more (**Fig. 4C**, **Fig. S7A**). Comparisons to the input samples collected at SynCom assembly, confirmed that all strains were present in the initial SynComs (**Fig. S7B**). While the composition of the metSC was more similar compared to the control and 500 µM MBOA treatments, several core strains of both SynComs differed in their relative abundances at high MBOA levels: *Stenotrophomonas* LST17 (susceptible to MBOA; tolerance from **Fig. 1A** and **S2**), which made up about 70% of the community in the DMSO and MBOA 500 µM treatment, was strongly reduced to 22% and 47% in the nonSC and metSC, respectively. In contrast, *Enterobacter* LMX9231 (intermediate tolerance) and *Pseudomonas* LMX9 (tolerant), as well as *Chitinophaga* LMN1 (susceptible) increased in their relative abundances at high MBOA concentration relative to the DMSO and 500 µM treatments. In conclusion, MBOA reduces bacterial community size (**Fig. 4A**), and influences community composition more strongly if a SynCom is unable to metabolize the compound (**Fig. 4B&C**).

## Discussion

Root-derived plant specialized metabolites structure microbiomes (9) and select root-associated bacteria that can both tolerate (11) and/or metabolize (40) these compounds. In this study we found that maize root bacteria, assembled as a synthetic community, interacted to metabolize benzoxazinoids *in vitro,* leading to the accumulation of the alternative MBOA- metabolization product HMPAA. The capacity to metabolize benzoxazinoids appeared to be beneficial to the metSC SynCom, enhancing the tolerance to MBOA and enabling the community to reach higher population densities than the nonSC SynCom at high MBOA concentrations. Furthermore, the effect of MBOA on community composition was dependent on the metabolization phenotype. Below we discuss the underlying mechanisms and benefits for microbial communities.

### Interacting bacteria redirect BX metabolism

Members of microbial communities often interact to degrade complex chemicals such as antimicrobial compounds, pollutants, and complex substrates but also cooperate for substrate utilization (24). Here, we found that SynComs of maize root bacteria interacted in the metabolization of MBOA. Against the expectation from our previous study, where we reported that single-strain cultures of *Microbacterium* LMB2 metabolized MBOA to AMPO (40), we found that the MBOA metabolizing metSC SynCom formed HMPAA as a dominant metabolite. Since none of the metSC strains formed HMPAA in single-strain culture, we concluded that HMPAA formation required the sole MBOA-degrading bacterium *Microbacterium* LMB2 in combination with at least one other strain. In paired cultures *Pseudomonas* LMX9 had the strongest effect of MBOA degradation and lead to complete AMPO absence, thus this strain appears to be the strongest contributor among the SynCom strains. Similar alternative product formation during benzoxazinoid degradation has previously been reported for a co-culture of the fungus *Fusarium verticillioides* with the bacterium *Bacillus mojavensis* (42). The *Fusarium* alone metabolized BOA to HMPA, but in co-culture with *Bacillus*, the intermediate AP was oxidized to the APO. Since the *Fusarium* is susceptible to APO while the *Bacillus* is not, this metabolic interaction resulted in inhibition of the pathogenic fungus without negatively affecting the bacterium. Here, we found that interacting bacteria redirected benzoxazinoid metabolization to an alternative compound.

### Mechanism of alternative MBOA metabolization

The formation of the alternative MBOA metabolization product HMPAA by the metSC SynCom required at least two strains and was only observed when *Microbacterium* LMB2 was present. Therefore, it is likely that LMB2 degraded MBOA by the previously identified lactonase BxdA (40) to the intermediate AMP, which is then acetylated by a second strain to HMPAA. There is evidence pointing towards such an acetylation as second step: *Pseudomonas chlororaphis* for example possesses an arylamine N-acetyltransferase (NAT1) that acetylates AP to AAP (47). It is possible that some SynCom strains possess homologues of NAT1 which analogously could act on AMP, a methoxylated AP, resulting in HMPAA formation. Further genomic and functional work with knockouts and transcriptomics is required to confirm the role of NAT1 homologs in HMPAA formation. Alternatively, a second pathway was proposed for the production of the acetamide AAP: acylation of AP to the malonylated form HMPA followed by deacetylation to AAP (48). Detoxification of benzoxazinoids via ring-opening and malonylation is well-known for fungal pathogens such as *Fusarium* species (41, 49). Throughout our analyses, we did not detect the malonylated intermediate HMPMA. However, as HPMA can be degraded by microorganisms such as Bacilli with and without the accumulation of AAP (48), the absence of HMPMA might also be explained by further degradation of such a product. Future work will focus on the identification and characterization of homologs of arylamine N-acyltransferases in the tested SynCom strains. *Pseudomonas* LMX9 will be especially interesting in this regard as it is the only strain that led to complete absence of AMPO in paired culture with *Microbacterium* LMB2 and also seemed to have the strongest effect on MBOA degradation in single dropouts of the metSC (**Fig. 2**). Our benzoxazinoid metabolizing SynCom thus offers a great tool to disentangle the biochemistry of bacterial interaction for metabolization of plant specialized metabolites.

### Interaction in benzoxazinoid metabolization is beneficial to community fitness

Benzoxazinoids were shown to selectively inhibit bacterial growth (11, 50), are metabolized by individual bacterial members of the maize root microbiome (40–42) and alter the composition of root microbial communities (14–16). Here we found that the metSC SynCom was more tolerant to MBOA than the nonSC SynCom (**Fig. 4A**). MBOA not only altered the size but also the composition of both SynComs implying that certain strains were more affected by the presence of MBOA than others (**Fig. 4C**). Specifically, high MBOA concentrations strongly reduced *Stenotrophomonas* LST17, which dominated the SynComs containing no or low amounts of MBOA, whereas *Pseudomonas* LMX9 and *Chitinophaga* LMN1 increased in relative abundance at the high MBOA concentrations. It is currently unclear, whether this increase in LMX9 and LMN1 was due to an increase in absolute abundance or whether the abundance of the strains remained stable and relative abundance solely increased due to the reduction of *Stenotrophomonas* LST17. For LST17, the inhibition was linked to its low tolerance to high concentrations of MBOA (**Fig. S2**). In the metSC SynCom, the *Stenotrophomonas* remained the most abundant strain, which could be explained by MBOA degradation and implies that this strain was also tolerant to the produced HMPAA.

It is currently unclear how the population dynamics developed over the course of the experiment. Even though *Microbacterium* LMB2 is required for MBOA metabolization in the metSC, it was only detected at low concentrations at the end of the experiment. One possible explanation is that LMB2 was present at higher concentrations at the beginning of the experiment, but then after initiating the MBOA metabolization, it was overgrown by fast-growing and less MBOA tolerant strains. In line with this, *Stenotrophomonas* LST17 would have been first present at low abundance and then started growing once MBOA concentrations fell below inhibitory concentrations. Interestingly, *Chitinophaga* LMN1 that also showed hardly any growth at 2’500 µM MBOA *in vitro* increased stronger in relative abundance in the nonSC than in the metSC at high MBOA. This suggested that other factors than MBOA tolerance contributed to the final SynCom composition such as microbe-microbe interactions based on metabolic exchanges between SynCom strains or modulation of direct competitive interactions (51, 52). Cross-feeding was for example previously identified in a 7-member community of maize root bacteria (53). An *Enterobacter* had previously been identified as keystone species in this SynCom (54). The keystone *Enterobacter* had the broadest substrate utilization of maize root extracts and its dropout decreased carbohydrate utilization of the SynCom (53). Our SynCom also contained an *Enterobacter* showing high abundance and substrate use may further explain community structure independent of tolerance to benzoxazinoids. Nevertheless, our observation that benzoxazinoids structured SynComs also *in vitro*, underlines the importance of benzoxazinoids in shaping root-associated microbial communities on benzoxazinoid producing maize roots (14–17).

### Effect of benzoxazinoid metabolization for bacterial growth and potential effects on host health

It remains an open question why microbes metabolize benzoxazinoids. Whether it is to detoxify them, to profit from a carbon source, to use degradation products as defence against other microbiota members or whether degradation is beneficial for the host. We found that a benzoxazinoid metabolizing SynCom was more tolerant to MBOA at high MBOA concentrations, which suggests specific adaption of host microbiomes to tolerate and metabolize host specialized metabolites and potentially presents a strategy for rhizosphere bacteria to access an extra carbon source to thrive in the rhizosphere (53, 55). Indeed, MBOA can be used as a sole carbon source by the degrading *Microbacterium* LMB2 (40). However, our SynCom experiments in complex medium suggest that under the tested conditions MBOA metabolization served mainly for detoxification for some microbiome members rather than strongly contributing to overall population size as a separate carbon source. As degradation of specialized metabolites can be highly context and environment dependent, the SynCom work could be next transferred to a gnotobiotic plant system as tested by Niu et al. (54). Members of the genus *Stenotrophomonas* show beneficial effects for plant growth and health (56). In a first step one could therefore focus on whether presence of MBOA metabolization increases local *Stenotrophomonas* abundance also *in planta* and potentially affects SynCom functions supplied to the host such as protection against plant pathogens. Alternatively, benzoxazinoid degradation and degradation products could additionally ineluence community function as regulators (57). Given the widespread nature of microbial interactions and the diversity of plant specialized metabolites, we propose that microbial interactions affect how microbial communities cope with plant specialized metabolites across the plant kingdom and thereby also potentially ineluence plant fitness.

## Conclusion

Natural microbial communities are diverse and single members interact to fulfil a complex task or to use resources efficiently (24). Here we report that maize root bacteria interact to tolerate and metabolize benzoxazinoids. Especially at high benzoxazinoid concentrations, the interaction of maize root bacteria positively affected the performance as a community. Our findings highlight the importance of studying plant specialized metabolites mediating chemical communication in plant-microbiome interactions in more complex synthetic or natural microbial communities.

## Materials and Methods

### Bacteria, SynCom assembly and experimental setup

Maize root bacteria (i.e., MRB collection) (11) were routinely grown on solid 100% TSA plates (30 g/L tryptic soy broth and 15 g/L of agar, both Sigma-Aldrich) or in liquid 50% TSB medium (15 g/L tryptic soy broth) with 180 rpm shaking at 25 – 28 °C. For cryo stocks, bacteria were grown for 48 h in liquid 100% TSB (30 g/L tryptic soy broth) and mixed with sterile-filtered glycerol (Sigma-Aldrich) at a final concentration of 20%.

The SynComs were assembled from taxonomically different MRB strains (11) that are distinguishable by 16S rRNA gene amplicon sequencing and differ in their ability to metabolize MBOA on their own (40). The SynComs were composed of the following six core strains, which do not metabolize MBOA: *Stenotrophomonas* LST17, *Pseudomonas* LMX9, *Enterobacter* LMX9231, *Chitinophaga* LMN1, *Bacillus* LBA21, *Streptomyces* LMG1. With the seventh strain, the SynComs differed in their ability to metabolize MBOA: *Microbacterium* LMI1x is unable to metabolize MBOA and was selected for the *non*-metabolizing SynCom (nonSC). *Microbacterium* LMB2 metabolizes MBOA on its own to AMPO and was part of the MBOA-*metabolizing* SynCom (metSC).

The individual strains were pre-grown in liquid cultures for the different experiments. For experiments depicted in Figs 1-3, pre-cultures were inoculated from single colonies and grown in 1 mL of liquid 50% TSB in 2 mL 96-well deep-well plates (Semadeni, Ostermundigen, Switzerland). Pre-culture plates were covered with a Breathe-Easy membrane (Diversified Biotech, Dedham, USA) and grown until stationary phase for 4 days at 28°C and 180 rpm.

For preparation of SynComs or strain mixes, cultures of individual strains were adjusted to OD_600_ = 0.6 and the required strains mixed at equal ratios. Main cultures were inoculated by adding 4 µL of the adjusted mixes/single strains into 96-well microtiter plates (Corning, Corning, USA) containing 200 µL of fresh liquid media including the compounds at the concentrations to be tested. The chemical treatments were prepared at the desired concentrations by mixing their stock solutions into the liquid growth media. The stock solution for MBOA (Sigma-Aldrich) was prepared in the solvent DMSO (Sigma-Aldrich) at 606 mM (100 mg/mL). MBOA was added to the medium at final concentrations of 0, 500 or 2’500 µM and the DMSO concentration was kept constant in all treatments including the controls.

We mainly utilized our previously described 96-well liquid culture based growth system (11). In brief, bacterial growth was monitored in 96-well plates in a high-throughput manner using a stacker (BioStack 4, Agilent Technologies, Santa Clara, USA), which was connected to a plate reader (Synergy H1, Agilent Technologies, Santa Clara, USA). This system shakes the cultures for 2 min (approx. every 80 minutes) before recording the OD_600_ readings in the plate reader. In each plate, wells with growth medium were included as no bacteria controls (NBC) and in each run one plate containing only media was included to monitor potential contaminations. For assessing community structure, precultures and SynComs were grown in Erlenmeyer elasks instead of 96-well plates and growth was assessed by OD_600_ measurements using a biophotometer (Eppendorf, Hamburg, Germany) and by dilution series plating and colony-forming units determination.

For details on the different experiments performed see **Supplementary Methods**.

### Data analysis

We used the R version 4.0 (R core Team, 2016) for all statistical analysis and visualization of the data. We first inspected the bacterial growth data for normality using Shapiro-Wilk tests. Differences between the two SynComs (nonSC vs. metSC) and treatments were assessed using pairwise t-tests and reported for the different treatments in the graphs. P-values were adjusted for multiple hypothesis testing using the Bonferroni method within R. We used the following packages for data analysis and visualizations: Tidyverse (58), Broom (59), DECIPHER (60), DESeq2 (61), emmeans (62), ggthemes (63), pheatmap (64), multcomp (65), phyloseq (66), phytools (67), vegan (68) in combination with custom functions. All source data and R code used for statistical analysis and graphing are available from https://github.com/PMI-Basel/Thoenen_et_al_SynCom.

### Metabolite analysis from bacterial cultures

Samples for metabolite analyses were processed and measured as described previously (40). Brieely, metabolites in bacterial cultures were fixed in 70% MeOH + 0.1% FA. Replicates of the same sample group were pooled and diluted followed by filtration and centrifugation to remove bacterial debris. Benzoxazinoids and their degradation products were profiled using an Acquity I-Class UHPLC system (Waters, Milford, USA) coupled to a Xevo G2-XS QTOF mass spectrometer (Waters, Milford, USA) equipped with a LockSpray dual electrospray ion source (Waters, Milford, USA). Different benzoxazinoid standard compounds were run together with the samples at varying concentrations and used for quantification and identification. Here in this work, we only report MBOA, AMPO and HMPAA. For details see Supplementary Methods. All source data and R code used for graphing are available from https://github.com/PMI-Basel/Thoenen_et_al_SynCom.

### Community analysis

DNA was extracted from 1 mL of SynCom samples (Experiment 4) using the NucleoSpin Soil kit (Macherey-Nagel, Düren, Germany) following the manufacturer’s instructions (see also **Supplementary Methods**). The 16S rRNA gene was amplified in a two-step PCR, with custom barcodes for sample identification added in the second PCR. The detailed method for PCR amplification and their primers, product purification and pooling is described in the **Supplementary Methods**. The pooled library was sequenced on a MiSeq instrument (Illumina Inc., San Diego, USA) with a v2 nano flow cell and the 2x250 bp pair-end sequencing protocol at the Next Generation Sequencing Platform (https://www.ngs.unibe.ch, University of Bern, Switzerland). The raw sequencing data was deposited at the European Nucleotide Archive (http://www.ebi.ac.uk/ena) under project number PRJEB86484.

The raw sequencing data were processed in R using the DADA2 pipeline version 1.20 (69). For details see **Supplementary Methods.** The 16S rRNA gene sequences of the SynCom members were mapped to the amplicon sequences using usearch (70) with an identity of 0.97. The counts of the mapped ASVs were normalized by rarefaction using phyloseq (66) and the abundances of the strains were further adjusted based on their estimated copy numbers of the 16S rRNA genes (**Dataset S1**). Compositional differences between the SynComs and by the treatments were assessed using Permutational Analysis of Variance (PERMANOVA, 99’999 permutations; model: ∼ SynCom * Treatment) on Bray Curtis distances using the R package vegan (68). The effects on community composition using the same model were visualized with a Canonical Analysis of Principal Coordinates (CAP) using the R package phyloseq (66). All source data and code used for statistical analysis and graphing are available from https://github.com/PMI-Basel/Thoenen_et_al_SynCom.

## Data availability

All source data is made publicly available. The sequencing data is available from the European Nucleotide Archive as Bioproject PRJEB86484. We provide all data on bacterial growth and metabolite data together with their analysis code on GitHub (https://github.com/PMI-Basel/Thoenen_et_al_SynCom).

## Code availability

All code used for statistical analysis and graphing is available from GitHub (https://github.com/PMI-Basel/Thoenen_et_al_SynCom).

## Acknowledgments

We thank Corinne Suter for support with culturing bacteria and plating assays and Mirco Hecht for supporting metabolomic analysis. Further, we thank Dr. Thomas Roder for the support with the open genome browser, Dr. Pamela Nicholson from the Next Generation Sequencing Platform in Bern for technical support with sequencing. This work was mainly supported by the Interfaculty Research Collaboration “One Health” of the University of Bern. It has also received support by grants of the European Research Council (No. 949595 to C.R.) and the Swiss National Science Foundation (No. 189071 to C.R.). Finally, the work was supported by the State Secretariat for Education, Research and Innovation SERI-funded ERC Consolidator Grant “mifeePs” (No. M822.00079 to K.S.).

## Supplementary Methods

### Experiments

#### Experiment 1 (Figure 1A and S2)

We cultured the nonSC and metSC SynComs in 96-well plates containing liquid 50% TSB supplemented with a final concentration of 500 µM MBOA. The experiment was stopped at 68 h. SynCom and NBC samples were grown in triplicates and pooled 1:1:1 for metabolite analysis. Fixing of bacterial cultures and metabolite analysis was performed as detailed below.

#### Experiment 2 (Figure 1B)

We repeated Experiment 1 with the nonSC and metSC SynComs (liquid 50% TSB, 500 µM MBOA) but as a time-series to characterise the kinetics of MBOA degradation and AMPO or HMPAA formation. For metabolite sampling, we removed replicate plates from the stacker at 16, 24, 44, 68 and 96 h. SynCom and NBC samples were grown in triplicates and pooled 1:1:1 for metabolite analysis. Fixing of bacterial cultures and metabolite analysis was performed as detailed below.

#### Experiment 3 (Figure 2)

We tested the nonSC and metSC SynComs, relative to single strain dropout SynComs (the metSC, minus the indicated strain) and each strain in paired cultures with *Microbacterium* LMB2. These assays were performed in liquid 50% TSB containing 500 µM MBOA and were stopped at 68 h. SynComs, the paired cultures and their non-bacterial controls (NBC, media with MBOA but without bacteria) were grown in triplicates and pooled 1:1:1 for metabolite analysis. Fixing of bacterial cultures and metabolite analysis was performed as detailed below. Classification of compound levels as “low” (>30% MBOA degraded compared to the NBC, <10% of max. AMPO-former (LMB2), <10% of max. HMPAA-former) or “high” (>90% MBOA degraded, >10% of max. AMPO-former (LMB2), >10% of max. HMPAA-former), were done as in our previous study (1).

#### Experiment 4 (Figure 3)

In parallel with Experiment 1, we tested the single strains and both complete SynComs, each with 5 replicates, for whether they could use MBOA as sole carbon source for growth. We followed the same procedure as described above but used minimal media (described previously (2)) containing MBOA at concentrations of 500 and 2’500 µM. As positive controls for growth, we grew the bacteria in minimal medium supplemented with glucose (500 and 2’500 µM) as sole carbon source. We continuously recorded the OD_600_ of the cultures and stopped the experiment at 68 h. For a measure of bacterial growth, we calculated the area under the growth curve (AUC, x-axis for time and y-axis for OD_600_) of the OD_600_ readings using the function *auc()* from package MESS (3) in R. We normalized growth in each treatment relative to the control. See analysis scripts for details.

#### Experiment 5 (Figure 4)

For scale reasons, this experiment was performed in Erlenmeyer flasks and not in 96-well plates as the other experiments. Modifications include that the strains were pre-cultured overnight in 50 ml Erlenmeyer flasks containing 30 ml liquid 50% TSB and then mixed in equal ratios (OD_600_ = 0.6) as nonSC and metSC SynComs. The assembled SynComs were pelleted (5 min at 3’600 rpm), washed twice with 10 mM MgCl_2_ buffer (Sigma-Aldrich, St. Louis, USA) and diluted in 35 mL of 10 mM MgCl_2_. Samples of these ‘input’ SynComs (i.e. SynComs at the start of the experiment when inoculating the Erlenmeyer flasks) were aliquoted in 1.5 ml microcentrifuge tubes (5 replicates) and stored at -80 °C for community analysis. The same concentration of 10 mM MgCl_2_ was added to the control treatment (NBC). Analogous to Experiments 1 – 4, cultures were grown in liquid 50% TSB supplemented with either 500 or 2’500 µM of MBOA or DMSO as a control. Five replicate cultures were set up for each of the 6 sample groups (2 SynCom * 3 treatments). The experiment was stopped at 68 h and the OD_600_ of each culture was recorded using a biophotometer (Eppendorf, Hamburg, Germany). Of each sample group, aliquots were fixed for metabolite analysis (details below). Metabolite measurements (n = 1) were made on pools of the five replicate cultures per sample group (pooled 1:1…). We also collected samples for community analysis by 16S rRNA gene amplicon sequencing (see below). For each of the 5 replicates per sample group, we collected two samples of the same culture resulting in n=10 replicates for community analysis. The samples were stored at -80 °C until processing. Finally, we assessed community size for each replicate (n=5) per sample group based on colony forming units (CFU) determined by plating serial dilutions on TSA plates and counting them at 22 °C.

### Metabolite analysis from bacterial cultures

Samples for metabolite analyses were processed and measured as described previously (1). To fix the metabolites in the bacterial cultures, we added 150 μL of the cultures to 350 μL of extraction buffer (100% methanol (MeOH) + 0.14% formic acid (FA)) in non-sterile round bottom 96-well plates (Thermo Fisher Scientific, Waltham, USA). We stored the fixed samples with a final concentration of 70% MeOH and 0.1% FA at -80 °C. To reduce the number of samples, we pooled replicates of the same sample group. We diluted the pooled sample by adding 50 to 700 μL MeOH 70% + 0.1% FA and filtered the cultures through regenerated cellulose membrane filters (CHROMAFIL RC, 0,2 µm; Macherey-Nagel, Düren, Germany) by centrifugation (3’220 g for 2 min) to remove bacterial debris. To pellet any residual particles, we centrifuged the extracts at 11’000 g for 10 min at 4 °C. We aliquoted the supernatants in analytical glass vials (Screw Neck Vials, 1 ml; VWR, Dietikon, Switzerland) and stored the samples at -20 °C until analysis.

We profiled the benzoxazinoids and their degradation products in the fixed and filtered extracts of the bacterial cultures using an Acquity I-Class UHPLC system (Waters, Milford, US) coupled to a Xevo G2-XS QTOF mass spectrometer (Waters, Milford, US) equipped with a LockSpray dual electrospray ion source (Waters, Milford, US). Gradient elution was performed on an Acquity BEH C18 column (2.1 x 100 mm i.d., 1.7 mm particle size; Waters, Milford, US) at 98– 50% A over 6 min, 50-100% B over 2 min, holding at 100% B for 2 min, re-equilibrating at 98% A for 2 min, where A = water + 0.1% FA and B = acetonitrile + 0.1% FA. The flow rate was 0.4 mL/min. The temperature of the column was maintained at 40 °C, and the injection volume was 1 μL. The QTOF MS was operated in sensitivity mode with a positive polarity. The data were acquired over an m/z range of 50–1’200 with scans of 0.1 s at a collision energy of 6 V (low energy) and a collision energy ramp from 10 to 30 V (high energy). The capillary and cone voltages were set to 2 kV and 20 V, respectively. The source temperature was maintained at 140°C, the desolvation temperature was 400 °C at 1’000 L/hr and the cone gas flow was 100 L/hr. Accurate mass measurements (<2 ppm) were obtained by infusing a solution of leucine encephalin at 200 ng/mL at a flow rate of 10 μL/min through the Lockspray probe (Waters, Milford, US). For each expected benzoxazinoid, standard compounds with four concentrations were run together with the samples (DIMBOA-Glc, DIMBOA, HMBOA, MBOA-Glc, MBOA, BOA, AMPO, APO, AAMPO, HMPMA, each at 10, 50, 200, and 400 ng/mL; HMPAA at 40, 200 ng/mL, 1 and 10 μg/mL). Here in this work, we only report MBOA, AMPO and HMPAA. The compound peaks in the raw chromatogram data were integrated using MassLynx 4.1 (Waters, Milford, US) and identified based on the reference compounds in the standards. Further, we searched for candidate compounds increasing in metSC over time using Progenesis QI Software (Waters, Milford, USA) and analysed it in R statistical software. All source data and R code used for graphing are available from https://github.com/PMI-Basel/Thoenen_et_al_SynCom.

### DNA extraction, library prep and sequencing

The DNA was extracted from two technical replicates per culture using the NucleoSpin Soil kit (Macherey-Nagel, Düren, Germany). For DNA extraction, 1 ml of culture was pelleted at 13’000 rpm, then the first buffer was directly added to the pellet. Afterwards the extraction was performed following the manufacturer’s instructions. The DNA concentration was quantified with the AccuClear® Ultra High Sensitivity dsDNA Quantitation Kit (Biotium, Fremont, USA). DNA was diluted to 0.2 ng/µl for subsequent amplification. A two-step PCR was performed, where with first PCR the DNA is amplified and in the second PCR the PCR products are tagged with custom barcodes. For the first PCR, the 16s rRNA gene was amplified with the specific primers 515-F (GTGYCAGCMGCCGCGGTAA, (4) and 806*-R (TTAGAWACCCBNGTAGTCC). The 806*-R was shortened by one base to avoid mismatch with the 16S sequence of the microbacteria strains. The PCR reaction mix was composed of 5 μl DNA template, 0.4 μl of 10 μM 515-F and 806*-R primers each, 8 μl 5Prime HotMasterMix (Quantabio, Beverly, USA), 2 μl 3% BSA and 4.6 μl autoclaved MilliQ water to a final reaction volume of 20 µl. The cycling profile was 94°C for 3 minutes, 25 cycles of 94°C for 45 seconds, 60°C for 60 seconds, 72°C for 90 seconds, and 72°C for ten minutes. All PCR reactions were purified with SPRIselect beads (Beckman Coulter Life Sciences, Indianapolis, USA) following manufacturer’s instructions. The reaction products from the first PCR were next amplified for another 10 cycles with uniquely barcoded primer pairs for each sample in a second PCR. The barcoded PCR products were purified with SPRIselect beads, DNA was quantified with the Qubit™ dsDNA BR kit (Invitrogen, Thermo Fisher Scientific, Waltham, MA, USA), and the samples were pooled equimolar using a Myra Liquid Handler (Bio Molecular Systems, Upper Coomera, Australia). The pooled library was bead purified and sequenced by MiSeq v2 500 cycle nano sequencing kit at the Next Generation Sequencing Platform (University of Bern) using the 2x 250 bp pair-end sequencing protocol (Illumina Inc., San Diego, USA).

### Community analysis details

#### The raw sequencing data was processed in R using the DADA2 pipeline version 1.20

(70). The analysis steps were wrapped in a Snakemake pipeline (5). In brief, raw reads were filtered and trimmed with the function *filterAndTrim()*. This step trims the 19 bp long primer sequences (trimLeft and trimRight = 19) and discards low-quality reads (truncQ = 2), reads matching PhiX and reads containing Ns. The reads were then subjected to the error learning step using *learnErrors()*, dereplicated using *derepFastq()* and denoised using dada(derepFs, err=errF, multithread=TRUE) following the DADA2 pipeline with default options. Next, the read pairs were merged using *mergePairs()* (minimal overlap of 20 bp) and chimera were removed using *removeBimeraDenovo()*. Finally, taxonomic assignment of the high-quality sequences was done against the SILVA database (v128; (6) using *assignTaxonomy()* and *addSpecies()* functions. A phyloseq object was exported for downstream analysis. **Dataset S2** documents all the dada2 pipeline. The source code for the analysis of the raw sequencing data is available from https://github.com/makrez/Analysis_documentation_Synthetic_communities_mrb.

For analysis of the community data, we first mapped the 16S rRNA gene sequences of the the SynCom members (all MRB strains have Sanger sequences available; (7) to the amplicon sequences using usearch (8) with an identity of 0.97. ASVs mapping to the SynCom members were used for calculation, other low abundant and non-mapping ASVs were discarded. The counts of the community data were normalized by rarefaction using phyloseq (9) and the abundances of the strains were further adjusted based on their estimated copy numbers of the 16S rRNA genes (**Dataset S1**). Compositional differences between the SynComs and by the treatments were tested with Permutational Analysis of Variance (PERMANOVA, 99999 permutations; model: ∼ SynCom * Treatment) on Bray Curtis distances using the R package vegan (10). The effects on community composition using the same model were visualized with a Canonical Analysis of Principal coordinates (CAP) using the R package phyloseq (9). All source data and code used for statistical analysis and graphing are available from https://github.com/PMI-Basel/Thoenen_et_al_SynCom.

## Supplementary Results

### Phenotype of *Enterobacter* LMX9231

One of the six core bacteria of both SynComs, *Enterobacter* LMX9231, had been identi∼ied earlier to colour MBOA-containing agar red, indicating AMPO formation (**Fig. S2A**, 1). Evaluating this strain in liquid culture, we found that, even though the strain forms AMPO on plates, it does not degrade MBOA to AMPO in the liquid growth condition (**Fig. S2B**). This ∼inding for liquid media was full consistent with all experiments of this study where the non-metabolising SynCom, of which LMX9231 is an abundant member (**Fig. 4C**), never degraded MBOA (see results of Experiments 1, 2 and 5). It appears plausible that *Enterobacter* LMX9231 may possess an alternative pathway for AMPO formation than Microbacteria and *Sphingobium*. When we found BxdA as an enzyme for the MBOA-to-AMPO metabolisation, we noticed that the best homologies to BxdA in *Enterobacteriaceae* showed less than 30% similarity on amino acid level. This low level of protein similarity was consistent with all tested strains that could not form AMPO-formation in liquid culture (**Fig. S2C**). We therefore think, that *Enterobacteriaceae* may possess another, BxdA- independent mechanism responsible for the AMPO-formation and this pathway seems to operate on Agar-plates and not in liquid cultures.

## Supplementary Figures

**Supplementary Figure S1:**
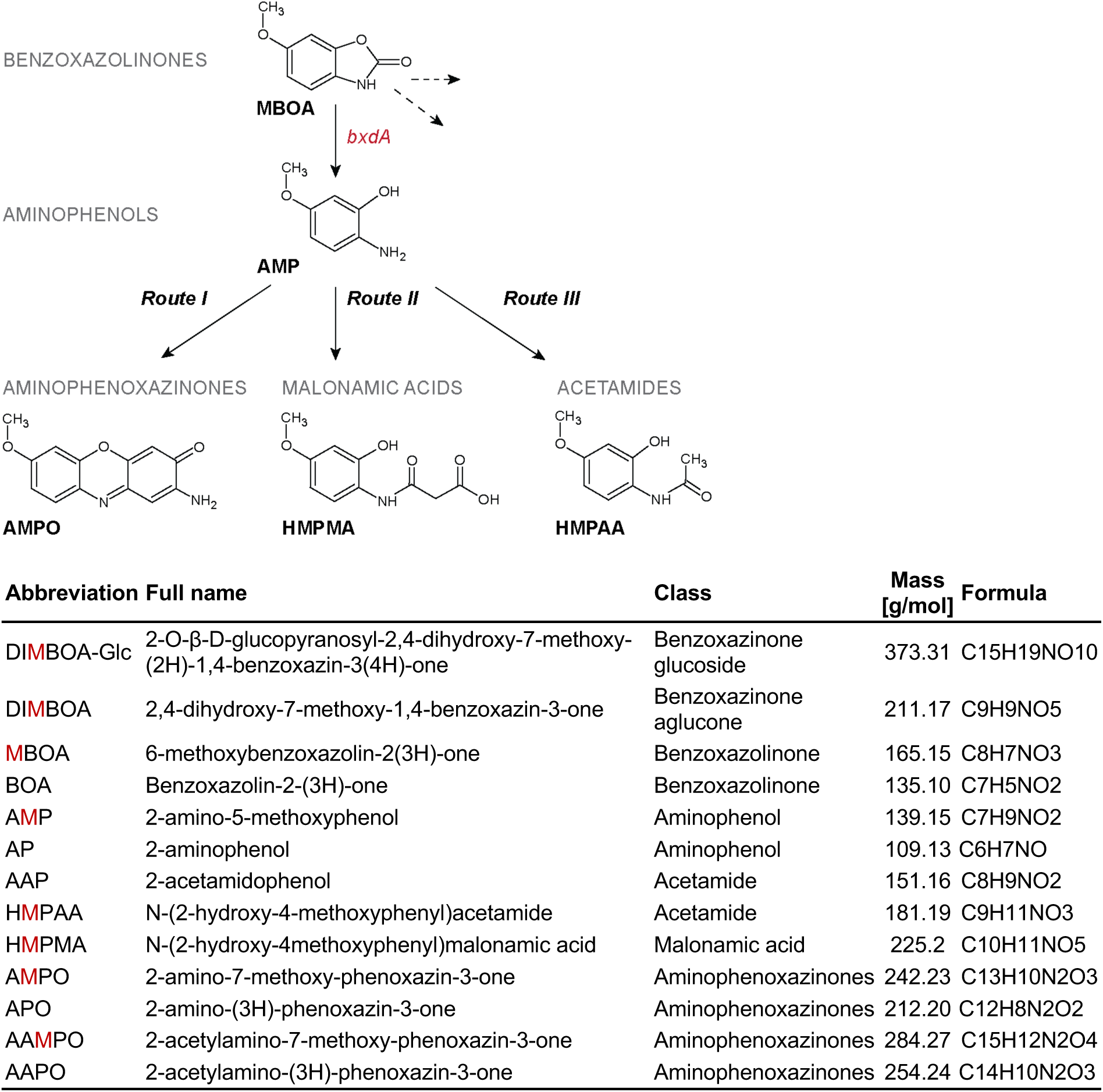
Benzoxazinoid metabolites produced by plants and degradation pathways in soil. The table lists the full chemical name, the compound class, the molar mass and the chemical formula.

**Supplementary Figure S2:**
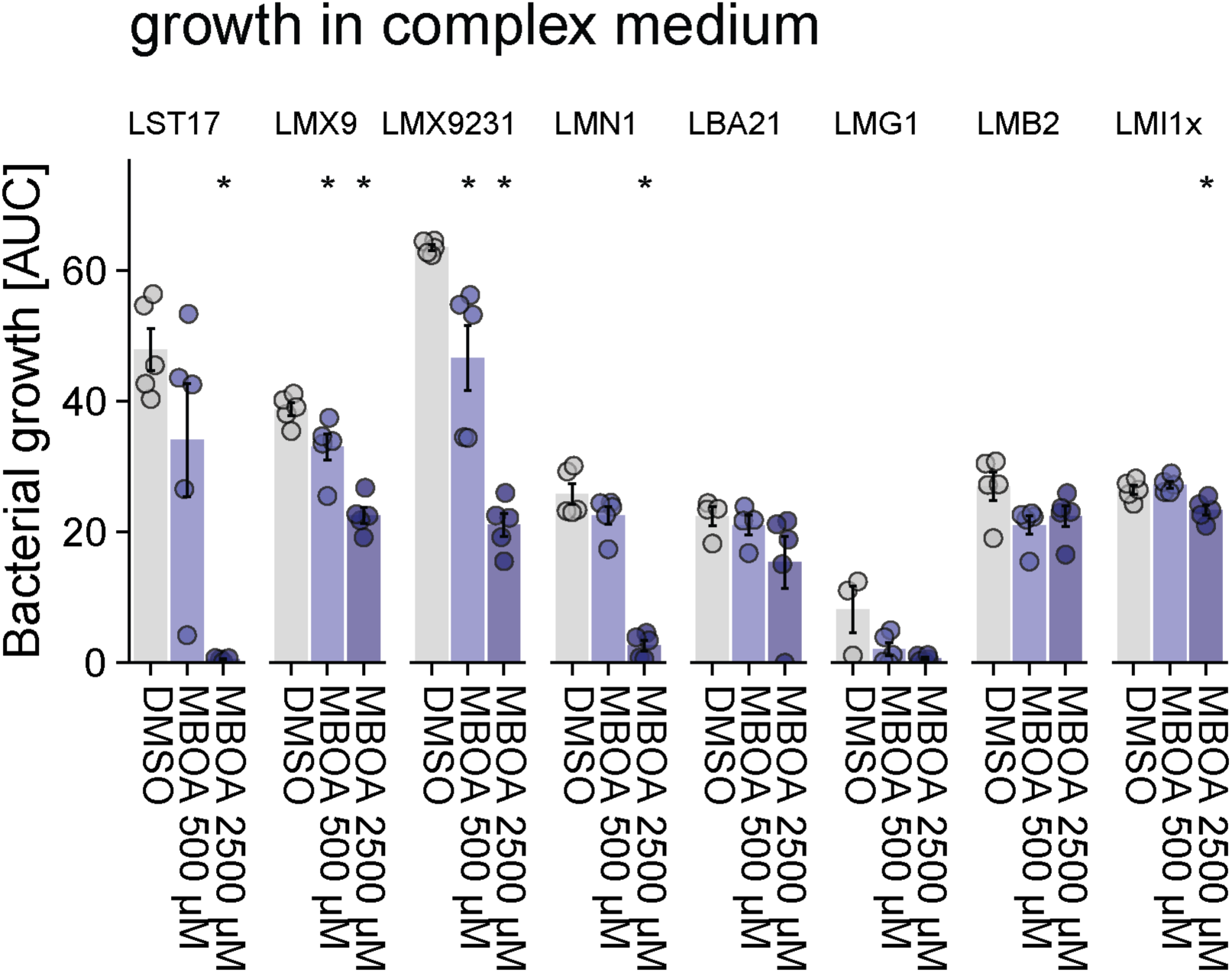
Growth of individual SynCom members in presence of MBOA. Bacterial growth was assessed in 50% TSB supplemented with MBOA (500 and 2’500 µM) or DMSO only (negative control). Five replicates were grown for each strain (n = 5). Means ± standard errors are reported and asterisks indicate significant differences between treatments and DMSO controls (pairwise t-test, Bonferroni-adjusted P < 0.05).

**Supplementary Figure S2:**
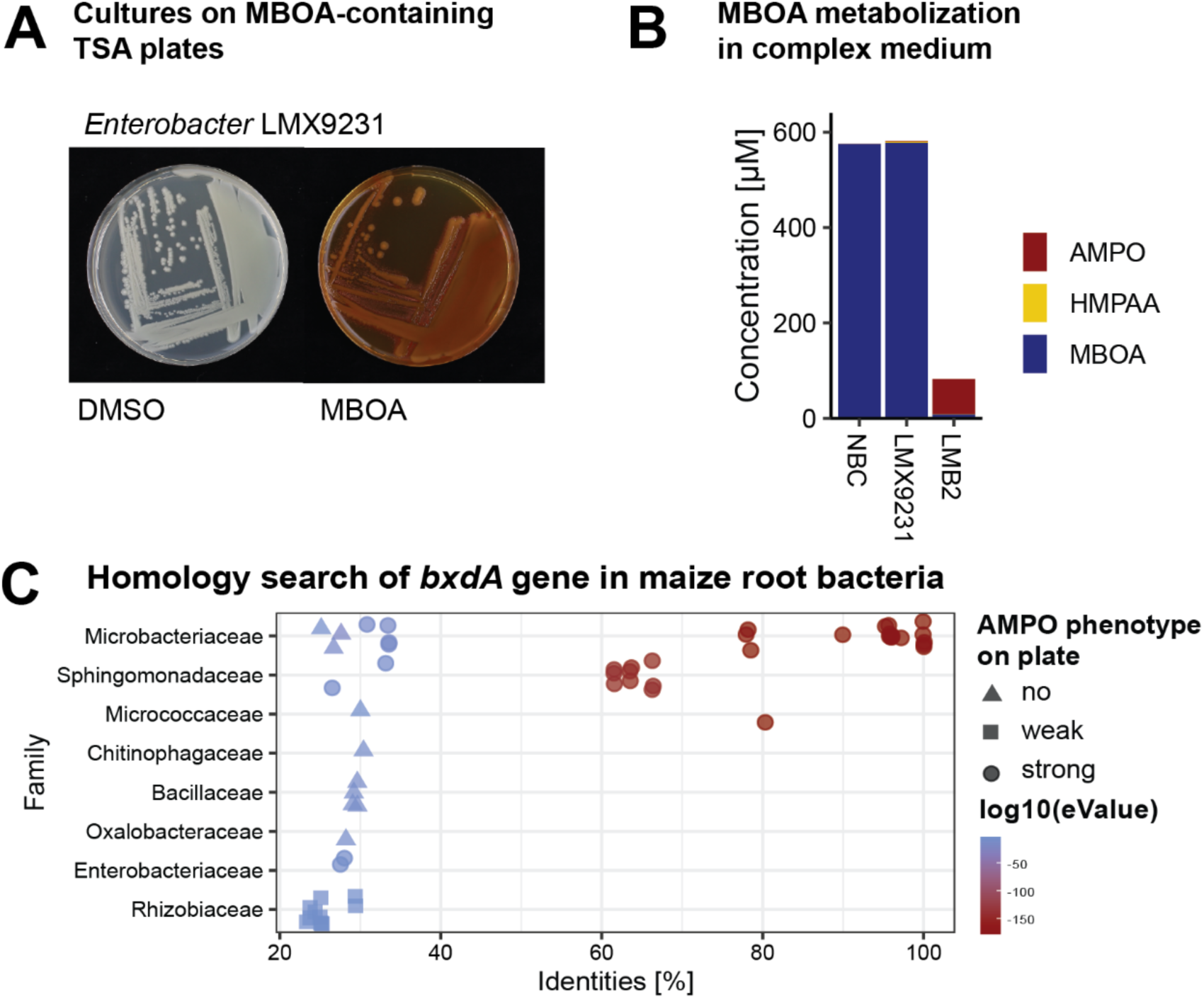
MBOA metabolisation phenotype of *Enterobacter* LMX9231: **A)** Pictures of *Enterobacter* LMX9231 grown on 100% TSA plates containing DMSO (2 mL/L) or MBOA (200 mg/L; ∼1’200 µM) for 10 days suggest AMPO formation. **B)** Metabolisation of MBOA and benzoxazinoid metabolites formed by *Enterobacter* LMX9231 and *Microbacterium* LMB2 (positive control) in 50% TSB supplemented with 500 µM MBOA show absence of MBOA metabolisation in liquid culture. **C)** Homology searches with the protein BxdA from *Microbacterium* LMB2 across all genome sequenced strains of the MRB collection. The panel C of this figure was shown in a previous publication (1).

**Supplementary Figure S3:**
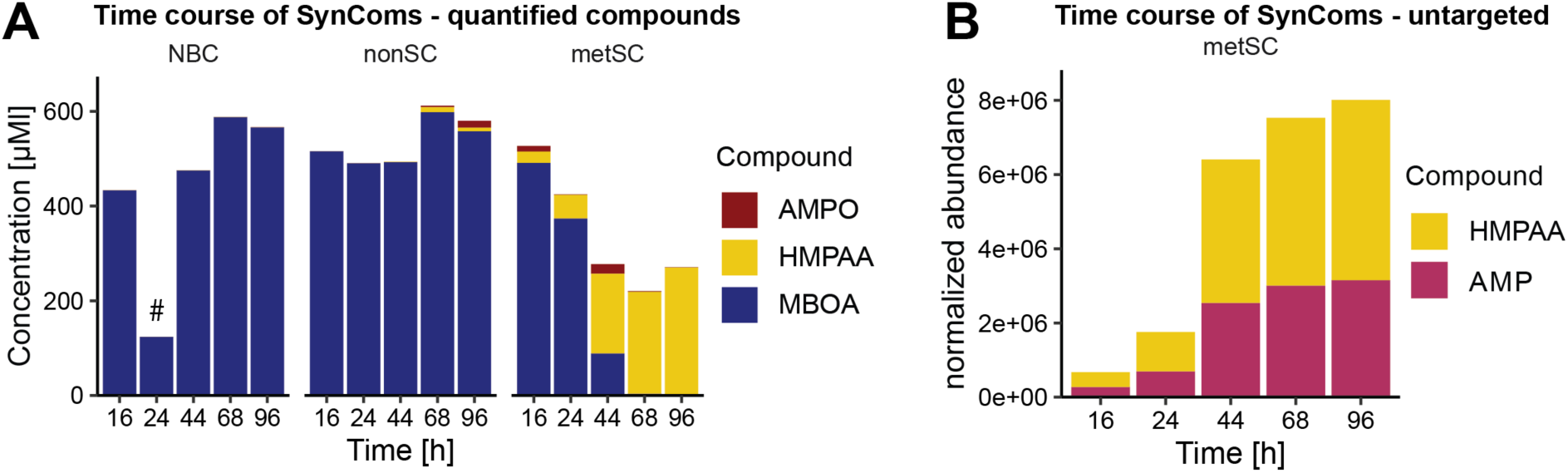
Metabolisation of MBOA by SynComs. The nonSC and metSC SynComs were grown in 96-well plates containing 50% TSB supplemented with 500 µM MBOA and samples were collected at the indicated time points up to 96 h. The 68 h time point is shown in Fig. 1B. **A)** The concentrations of MBOA, AMPO and HMPAA metabolites were quantified using analytical standards while in **B)** the intermediate AMP (see **Fig. S1**) is reported based on the normalized peak area (peak corresponding to the mass of AMP). HMPAA is reported again but on its normalized peak area for reference. Metabolite measurements (n = 1) were made on pools of three independently grown cultures (#: sample with failed pooling). Part of panel A of this figure (NBC) was shown in a previous publication (1).

**Supplementary Figure S4:**
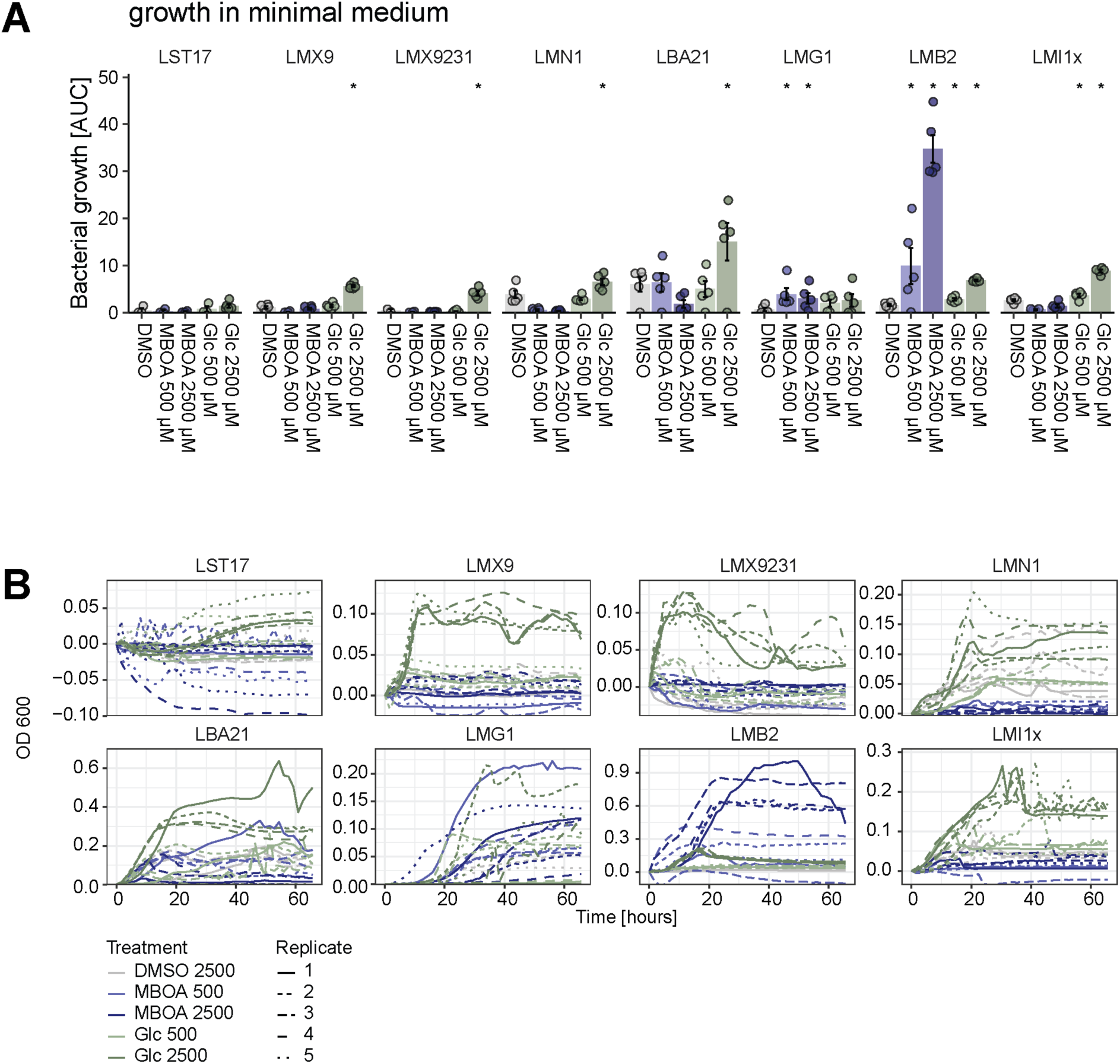
Growth of individual SynCom members in minimal media. Bacterial growth in the minimal medium with DMSO (negative control), MBOA (500 and 2’500 µM) or glucose (positive control; same concentrations) as sole carbon sources. Five replicates were grown for each strain (n=5). Means ± standard errors are reported and asterisks indicate significant differences against their DMSO control (pairwise t-test, Bonferroni-adjusted P < 0.05). Part of the data of this figure was shown in a previous publication (1).

**Supplementary Figure S5:**
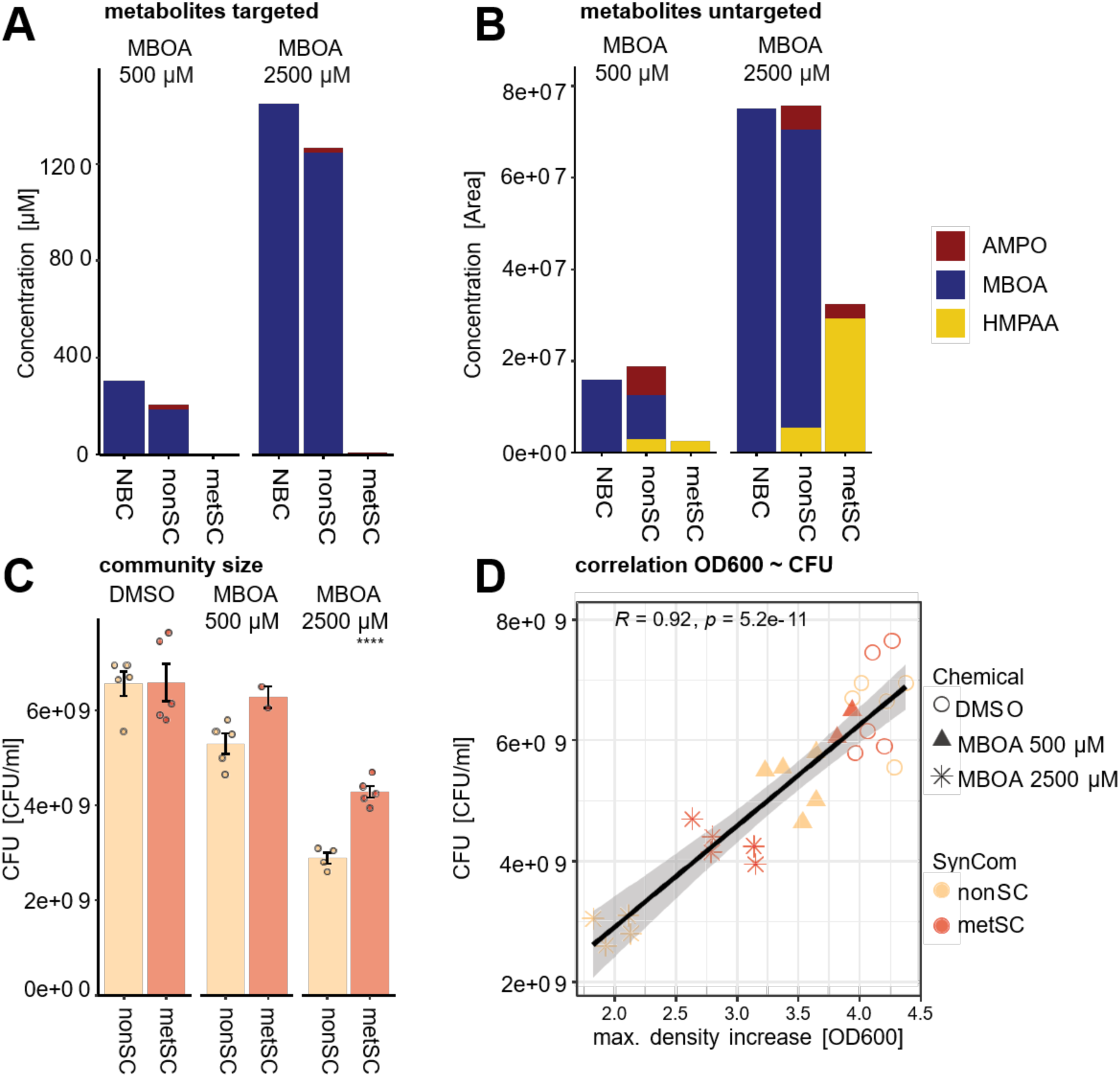
Metabolites and growth of the SynComs in presence and absence of MBOA. SynComs were grown in shake flasks containing 20 mL 50% TSB supplemented with DMSO or 500 µM or 2’500 µM MBOA until harvest at 68 h. Five replicate cultures were set up for each SynCom and treatment. **A)** Measurements of metabolites after exposure of both SynComs and their no bacteria control (NBC, media with MBOA but without bacteria) to MBOA. Metabolite measurements (n = 1) were made on pools of five replicate cultures. Note that HMPAA is not shown in A because of absence of a standard. **B)** Untargeted measurements of metabolites, relative concentrations were calculated from the area of the integrated peaks. **C)** Community size was determined by counting the colony forming units (CFU/mL) at the end of the experiment (n=5). Asterisks indicate significant differences between the SynComs (pairwise t-test, Bonferroni-adjusted P < 0.05). **D)** Corrleation of the two measurements for bacterial growth, OD_600_ compared to CFU counts (shown in Fig. 4A).

**Supplementary Figure 6:**
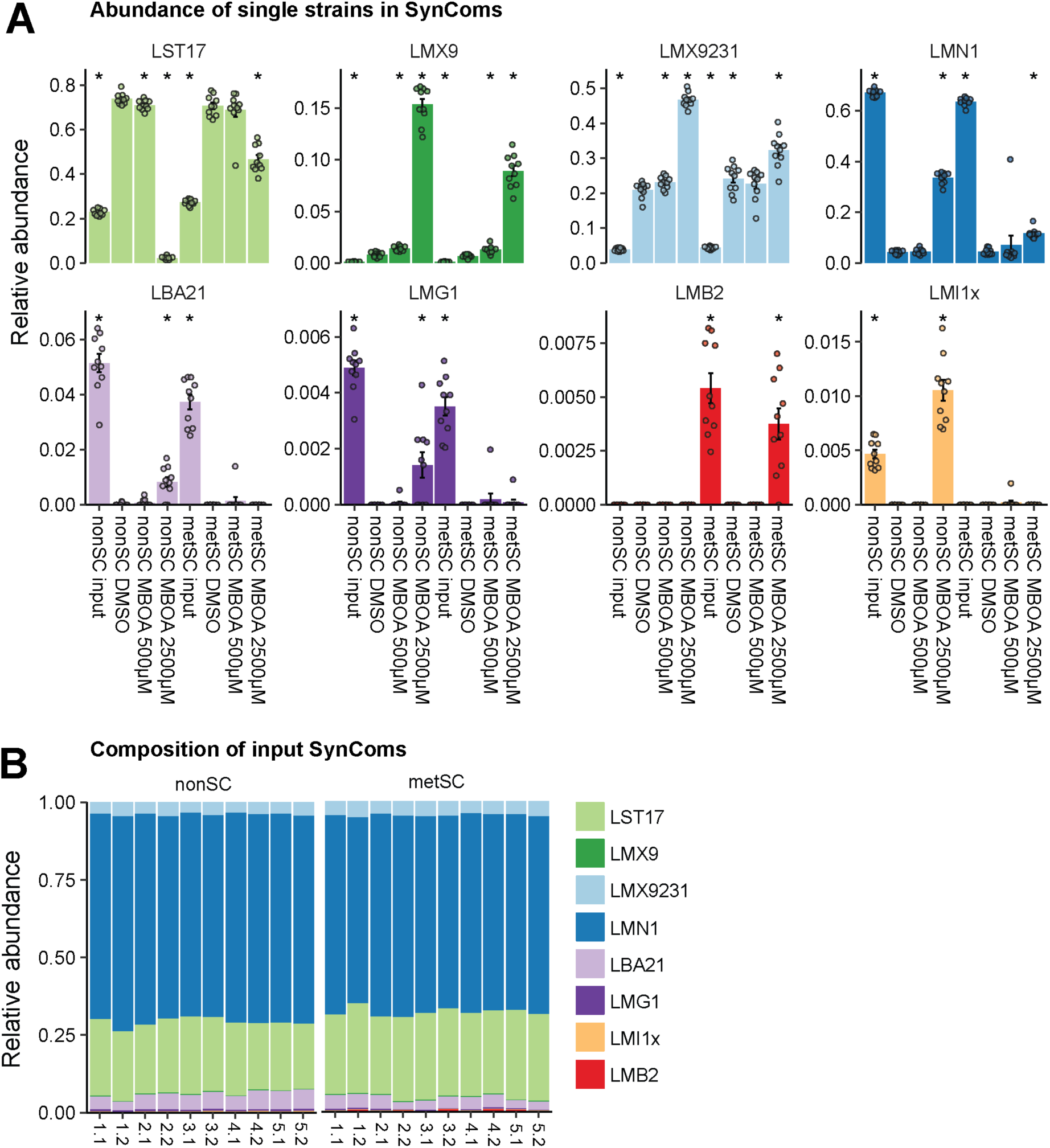
Abundance of single strains in the SynComs in presence and absence of MBOA. SynComs were grown in shake flasks containing 20 mL 50% TSB supplemented with DMSO or 500 µM or 2500 µM MBOA until harvest at 68 h. Five replicate cultures were set up for each SynCom and treatment and two parallel samples of the same culture were collected for community analysis by 16S rRNA gene amplicon sequencing. **A)** Relative abundances of each single strain in the SynCom at the beginning of the experiment (input) and treatment combinations during the experiment. Asterisks indicate significant differences between the treatments (pairwise t-test, Bonferroni-adjusted P < 0.05). **B)** Community analysis was also conducted with the ‘input’ SynComs, i.e. to determine the strain abundances at the start of the experiment.

## Supplementary Tables

**Table S1:** PERMANOVA of SynCom composition. Factorial effects on community compositions were assessed using PERMANOVA based on Bray Curtis distances and the model ∼SynCom * Treatment (n=10).

|  | **Df** | **SumOfSqs** | **R2** | **F** | **Pr(>F)** |
| --- | --- | --- | --- | --- | --- |
| **Treatment** | 2 | 0.7837848 | 0.2598361 | 10.467239 | **0.001** |
| **SynCom** | 1 | 0.0504273 | 0.0167174 | 1.346886 | 0.26 |
| **Treatment:SynCom** | 2 | 0.1604922 | 0.0532055 | 2.14333 | 0.063 |
| **Residual** | 54 | 2.0217547 | 0.6702411 | NA | NA |
| **Total** | 59 | 3.0164589 | 1 | NA | NA |

## Supplementary Datasets

**Dataset S1: Documentation of dada2 pipeline script used**. This file is a html markdown of the R script containing the code to process the raw sequencing reads from the community profiling (Experiment 5) to the phyloseq object which was then further analysed in R.

**Dataset S2:** Table contains information on all bacterial strains used in the SynCom, taxonomy information, 16S sanger sequence and the number of 16S copies.

